# Pervasive population genomic consequences of genome duplication in *Arabidopsis arenosa*

**DOI:** 10.1101/411041

**Authors:** Patrick Monnahan, Filip Kolář, Pierre Baduel, Christian Sailer, Jordan Koch, Robert Horvath, Benjamin Laenen, Roswitha Schmickl, Pirita Paajanen, Gabriela Šrámková, Magdalena Bohutínská, Brian Arnold, Caroline M. Weisman, Karol Marhold, Tanja Slotte, Kirsten Bomblies, Levi Yant

## Abstract

Ploidy-variable species allow direct inference of the effects of chromosome copy number on fundamental evolutionary processes. While an abundance of theoretical work suggests polyploidy should leave distinct population genomic signatures, empirical data remains sparse. We sequenced ∼300 individuals from 39 populations of *Arabidopsis arenosa*, a naturally diploid-autotetraploid species. We find the impacts of polyploidy on population genomic processes are subtle yet pervasive, including reduced efficiency on linked and purifying selection as well as rampant gene flow from diploids. Initial masking of deleterious mutations, faster rates of nucleotide substitution, and interploidy introgression all conspire to shape the evolutionary potential of polyploids.

## Introduction

Whole genome duplication events (i.e. polyploidizations) have occurred throughout the tree of life [1, 2] and are associated with biological phenomena of great socio-economic importance ranging from crop domestication [3] to cancer development [4]. The effects of polyploidy are far-reaching, ranging from single-cell level processes [5] through organism-level phenotypes [6], up to population genetic processes and biotic interactions in the ecosystem [7-9].

Polyploidy has enjoyed keen interest in population genetics, where the theoretical effects of chromosome copy number have been widely explored [10-14]. Theory predicts substantive differences between diploids and tetraploids for both neutral and selective processes [15, 16]. In autotetraploids (i.e. resulting from within-species genome duplication), levels of neutral polymorphism and diversity are expected to be doubled, and neutral divergence due to genetic drift should occur at half the rate expected in diploid populations [17]. Similarly, all else being equal, a doubling of the population-scaled recombination rate should reduce linkage disequilibrium in autotetraploids. Genome duplication is also expected to affect various types of selection, due to differences in the manifestation of allelic dominance in different ploidies. Equilibrium frequencies at mutation-selection balance can be orders of magnitude higher for recessive mutations in tetraploids, resulting in increased genetic load [18]. For selection on beneficial alleles, allele frequencies will change slower in higher ploidies for most patterns of dominance [19], although this can be more than compensated for by the doubled rate at which beneficial mutations are introduced in tetraploids [16, 20]. Lastly, weaker linkage in autopolyploids may reduce interference between beneficial alleles allowing greater opportunity for a beneficial allele to recombine onto haplotypes with fewer deleterious mutations.

In addition to the effects of polyploidy on genome evolution, another effect may come from an altered potential for gene flow with co-occurring diploid populations. Although polyploidization is traditionally viewed as a means of instant speciation because diploid/tetraploid hybridization is expected to result in low-fertility triploids [21-23], it has been shown that in at least some cases the ploidy barrier is permeable, particularly from diploids to polyploids [24, 25]. As the range of a polyploid lineage expands, it may encounter locally-adapted diploids, in which case interploidy introgression could supply genetic variation facilitating rapid local adaptation of the polyploid. Such adaptive introgression is increasingly being recognized as an important force in diploid systems [26, 27], yet genomic evidence in a ploidy variable system is lacking. Additionally, polyploidy may break down systems of reproductive isolation present in diploid progenitors. For example, although reproductive isolation between diploid *Arabidopsis arenosa* and *Arabidopsis lyrata* is near complete, tetraploid *A. lyrata* forms viable hybrids with both diploid and tetraploid *A. arenosa* [28].

However, the majority of theoretical expectations on population genomic effects of polyploidy remain untested at the genome level in natural polyploid systems. Studies of natural systems are invaluable for testing and further calibrating theoretical expectations, and numerous arguments, up to its role in carcinogenesis [4], call for better understanding of genome evolution in naturally evolving polyploid systems. Lack of empirical population genomic data is particularly pronounced for autopolyploids, which arise from within-species genome duplication and thus carry four equally homologous copies of each chromosome [29]. In contrast to the better studied allopolyploids, in which the effects of polyploidy are confounded with subgenome divergence, autopolyploids allow us to directly study the effects of chromosome number *per se*. Using a new model for autopolyploidy, *Arabidopsis arenosa* [30], we generated the most comprehensive genomic dataset to date of a natural autopolyploid and its diploid sister lineages and test theoretical predictions of genome duplication in a naturally evolving system.

*Arabidopsis arenosa*, the only representative of the model genus with both diploids and widespread autotetraploids, provides a powerful system for the study of the population genomic effects of genome doubling in nature. The tetraploids, tracing to a single origin tens of thousands years ago in the Western Carpathians, have spread across much of Europe, creating multiple contact zones with divergent diploid lineages [31, 32]. We present a range-wide analysis of 39 *A. arenosa* populations (287 resequenced genomes, 105 diploids from 15 populations and 182 tetraploids from 24 populations; Figure 1). We focus on four main questions concerning the genomic impact of selection and migration in a ploidy variable system: First, we investigate if purifying selection is relaxed in autotetraploids. Second, we ask whether the strength of linked selection is more pronounced in one ploidy versus the other. Third, we evaluate whether ploidy may alter rates of adaptation. Lastly, we look for evidence of interploidy gene flow at two independent contact zones to assess the impact of interploidy introgression on polyploid evolution. Overall, our empirical analyses provide novel insights into the complexity of autopolyploid evolution, supporting some but not all theoretical predictions. In tetraploids, altered selective processes as well as introgression alter the genomic landscape relative to diploids, possibly reshaping the evolutionary potential of the polyploid lineage. Such interacting features will likely apply generally to naturally evolving autopolyploid systems.

**Fig. 1.**
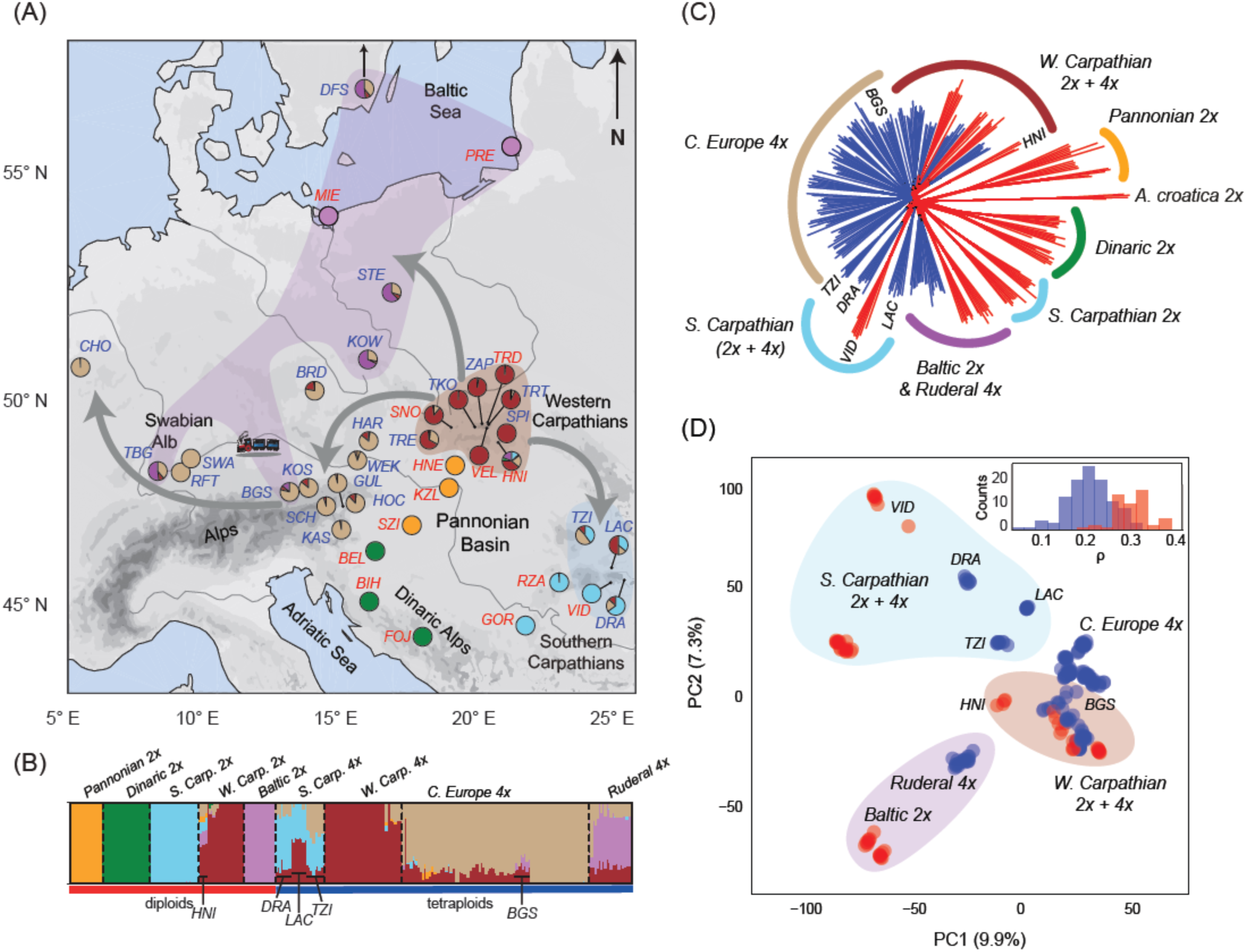
Geographic distribution and range-wide genetic variation of *Arabidopsis arenosa*. (A) Distribution of the 39 *A. arenosa* populations coloured according to major genetic groups inferred by Bayesian clustering based on fourfold degenerate SNPs (pie charts reflect proportion of individual cluster membership under K=6; red labels - diploid, blue - tetraploid populations). Arrows mark spread of distinct tetraploid lineages from the putative ancestral area in the W. Carpathians. (B) Posterior probabilities of cluster assignment of the 287 *A. arenosa* individuals. (C) Neighbor-joining network based on Nei’s genetic distances among all individuals including the outgroup *Arabidopsis croatica*. (D) First two principal components of all but the two most divergent diploid (*Pannonian* and *Dinaric*) *A. arenosa* lineages. The inset depicts genetic divergence (*ρ*) within each ploidy.

## Results

### *High diversity and population differentiation in natural* A. arenosa

*Arabidopsis arenosa* is an obligate outcrosser and all populations exhibit high genome-wide diversity (average pairwise *θ_π_* = 0.015, Table 1), an order of magnitude higher than that reported for the predominantly self-fertilizing *A. thaliana* [33]. All else being equal, polyploidy is expected to increase diversity due to increased effective population size (8*Neμ* in tetraploids versus 4*Neμ* in diploids). Although tetraploid populations exhibit slightly higher *θ*_*W*_ (Watterson’s theta) at non-synonymous 0-fold degenerate sites (0-dg), where any mutation results in an amino-acid change, we observe no significant increase of *θ_π_* or *θ*_*W*_ in tetraploid populations at putatively neutral 4-fold degenerate sites (4-dg), where no mutation results in an amino-acid change. However, we find a highly significant difference between ploidies for the ratio of 0-dg *θ*_*W*_ to 4-dg *θ*_*W*_ (p < 0.001), suggesting an additional role of selection in patterning tetraploid diversity (Table 1, S1-2). The impact of genome duplication on *θ*_*W*_ (in contrast to *θ_π_*) is consistent with tetraploid recent origin, as *θ*_*W*_ is more sensitive to the accumulation of rare variants.

**Table 1.** Measures of within-population diversity and among-population divergence in diploid and tetraploid *A. arenosa*

|  | Divergence |  |  | Diversity<sup>1</sup> |  |  |  |  |  |  |  |  |  |  |
| --- | --- | --- | --- | --- | --- | --- | --- | --- | --- | --- | --- | --- | --- | --- |
|  | Rho /<br>F<sub>st</sub><sup>2</sup> | AMO<br>VA<sup>3</sup> | rM<sup>4</sup> | pairwise diversity (θ<sub>π</sub>) |  |  | Watterson's θ (θ<sub>w</sub>) |  |  | Tajima's D |  |  | π<sub>NS</sub>/π<sub>S</sub> | θ<sub>NS</sub>/θ<sub>S</sub> |
| Sites | 4-dg | 4-dg | 4-dg | all | 4-dg | NS (0-dg) | all | 4-dg | NS (0-dg) | all | 4-dg | NS (0-dg) | - | - |
| Diploids | 0.30 / |  | 0.14 | 0.016 | 0.022 | 0.0054 | 0.015 | 0.022 | 0.005 | 0.03 | 0.16 | -0.09 | 0.242 | 0.255 |
| (14 pops) | 0.29 | 71 | n.s. | (0.003) | (0.003) | (0.0007) | (0.004) | (0.003) | (0.0009) | (0.21) | (0.18) | (0.23) | (0.017) | (0.017) |
| Tetraploids | 0.20 / |  | 0.55 | 0.015 | 0.023 | 0.0055 | 0.016 | 0.023 | 0.006 | -0.23 | 0.00 | -0.41 | 0.237 | 0.263 |
| (22 pops) | 0.11 | 48 | *** | (0.004) | (0.006) | (0.0013) | (0.004) | (0.005) | (0.0013) | (0.29) | (0.27) | (0.28) | (0.007) | (0.007) |
| Difference<sup>5</sup> | - | - | - | n.s. | n.s. | n.s. | n.s. | n.s. | . | ** | . | *** | n.s. | *** |

Populations with < 5 individuals were excluded; for populations with > 5 individuals, sites were randomly downsampled to five to facilitate comparison across populations.
<sup>1</sup>values averaged across populations within the ploidy, standard deviation is in parentheses
<sup>2</sup>values averaged over pairwise comparisons of populations belonging to that ploidy
<sup>3</sup>% of explained variance among populations (compared to variance within populations) in Analysis of Molecular Variance (AMOVA)
<sup>4</sup> Isolation by distance tested by Mantel test; the rM for diploid populations became 0.23\* when spatially distant but genetically proximal Baltic populations were excluded
<sup>5</sup> Wilcoxon rank sum test; n.s. non significant, $p \leq 0.07$ \* $p \leq 0.05$ , \*\* $p \leq 0.01$ , \*\*\* $p \leq 0.001$

At equilibrium, divergence due to genetic drift in tetraploids is expected to be half that in diploids [17]. In line with this (and the greater age of diploids), we find lower differentiation between tetraploid populations (Table 1, S3, Fig. 1D). The diploid populations form 5 divergent geographically separated groups, consistent with a previous restriction site associated DNA sequencing (RADseq) study [34], hereafter referred to as the *Pannonian, Dinaric, Baltic*, Southern Carpathian (*S. Carp.*), and Western Carpathian (*W. Carp*.) lineages (Fig. 1). While the *Dinaric-2x* and *Pannonian-2x* lineages were highly distinct (average pairwise F_ST_ from other diploid populations is 0.34 and 0.31, respectively), the two Carpathian diploid lineages (*S. Carp.-2x* and *W. Carp.-2x*) were less differentiated from each other (avg. F_ST_ = 0.25), consistent with occasional hybridization between Carpathian diploids in the past (*Baltic-2x* lineage, Table S4) and recently (HNI population, Fig. 1B). The tetraploids are split into four lineages that roughly correspond to the following geographic regions: Southern Carpathians (*S. Carp.*), Western Carpathians (*W. Carp.*), and the Alps together with western Central Europe (*C. Europe*). The fourth, *Ruderal,* group is the most widespread yet ecologically distinct, occupying man-made sites (e.g. railway ballast) from southern Germany to Sweden (Fig. 1). Groupings were consistent across a range of algorithms (Fig. 1, S1 – S5), although some methods identified finer sub-structure within the *S. Carp.-2x* and *C. Europe-4x* lineages (Fig. S5).

### Ploidy effects on purifying selection

Ploidy differences in patterns of diversity (Table 1) hint at relaxed purifying selection in tetraploids in line with theory [18]. To test this hypothesis, we compared gene level diversity with gene expression, which frequently correlates with selective constraints (e.g. [35-37]). We confirmed that highly-expressed genes exhibit reduced nonsynonymous diversity (significant effect of expression on *θ*_*W*_ at 0-dg sites as well as on the 0-dg/4-dg ratio of *θ*_*W*_; Fig. 2A, 2B and Table S5). This is the case in both ploidies, but we observed an overall significant increase in 0-dg/4-dg *θ*_*W*_ ratio in tetraploids driven by an increase of nonsynonymous diversity (Table 1, Fig. 2B, S6), which remained significant even when we included the estimated population size (number of haploid genomes, N_g_) in the model (Fig. 2B, Table S6). This confirms that beyond the increased mutational input resulting from the doubling of genome copies in tetraploids (doubled N_g_ for equal population sizes), there is an additional increase of non-synonymous diversity in tetraploids, likely from a relaxation of purifying selection. This increase affects all genes similarly across expression levels suggesting it is independent of functional constraints.

**Fig 2:**
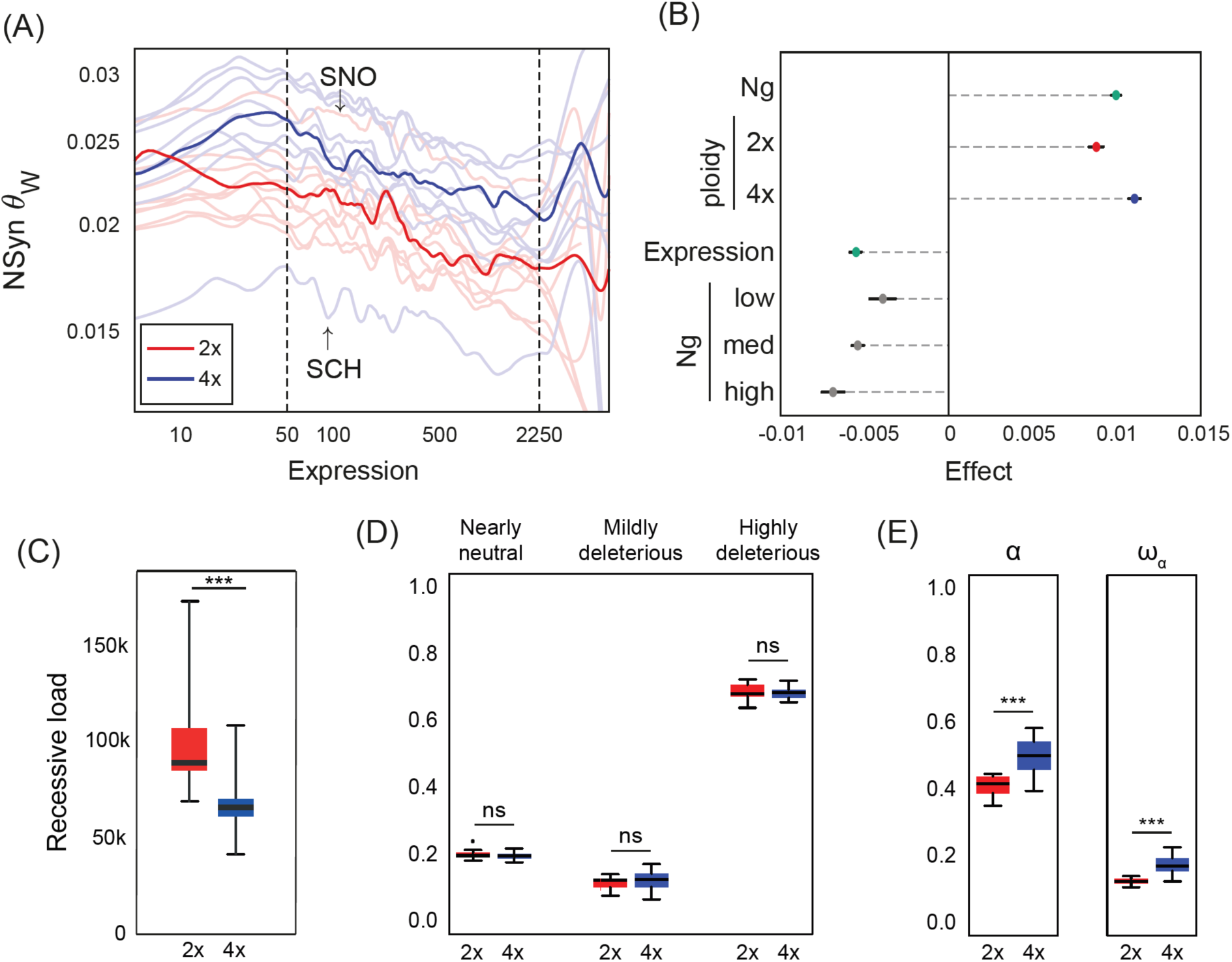
Effects of ploidy on purifying selection, genetic load, and the distribution of fitness effects (DFE). (A) Relationship of genic nonsynonymous diversity against relative log-expression shown for each population and ploidy (resp. faint and bold). (B) Estimated effects of haploid effective population size (N_g_) and levels of expression on nonsynonymous (0-dg) θ_W_ and their interaction terms with ploidy and N_g_ respectively in a multiple linear model. (C) Lower recessive load in tetraploid individuals estimated as number of 0-dg sites homozygous for a derived allele. (D) DFE in populations grouped by ploidy and binned according increasing strength of purifying selection. (E) Proportion of adaptive substitution (*α*) and proportion of adaptive substitution relative to neutral (*ω*_*α*_) of all populations grouped by ploidy.

Such relaxed purifying selection could be due to either a reduction in the strength of selection *per se* or simply because selection is less efficient. At a given allele frequency, homozygotes are much less frequent in tetraploid populations (*q*^*2*^ versus *q*^*4*^), and if mutations are recessive, the deleterious phenotype is rarely observed. This makes purifying selection inefficient relative to diploids even if the fitness costs of mutant homozygotes (i.e. selection strength) are equivalent across ploidies. To distinguish between these causes of reduced purifying selection, we evaluated the distribution of fitness effects (DFE) across both ploidies and find no apparent differences in the strength of purifying selection in diploid vs. tetraploid populations (Fig. 2D). Thus, purifying selection is not weaker *per se*, it is simply less efficient at reducing allele frequencies because deleterious mutations are better masked in autotetraploids.

That said, two assumptions in the DFE estimation method we use [38] deserve consideration. First, the method assumes a diploid model of mutation-selection-drift balance. Since allele frequencies at mutation-selection balance are expected to be higher in autotetraploids [15], the diploid model would be biased towards inferring weaker selection than necessary to explain the polyploid data. Second, all deleterious mutations are assumed to have additive effects on fitness. If deleterious mutations are recessive, equilibrium allele frequencies at mutation-selection balance can be orders of magnitude greater in tetraploids, which would amplify even further the first bias towards inferring weaker fitness effects in tetraploids. If purifying selection were truly weaker in tetraploids, these biases would make this more apparent; instead, we find no evidence for ploidy differences in the DFE (Fig. 2D, S7 and Table S7).

In the long run, the combined effects of inefficient purifying selection and increased mutational input in tetraploids is expected to allow for both higher numbers and higher frequencies of deleterious alleles, and consequently a higher genetic load (i.e. the average reduction in fitness of an individual relative to an optimal genotype bearing no deleterious alleles) [18]. However, tetraploid *A. arenosa* lineages may not have reached their new mutation-selection equilibrium given their relatively young age [31] and the gene flow they experience from diploids (discussed below). We therefore estimated genetic load, assuming recessivity of deleterious alleles, in both diploids and tetraploids by counting the per-individual number of homozygous genotypes for derived, nonsynonymous alleles in each population. By this measure, load is significantly lower in tetraploids than in diploids (Wilcoxon rank-sum test of population means, W = 264, p < 0.0001, Fig. 2C). It should be noted, however, that under a given dominance model, comparisons between ploidies are only strictly valid if the distribution of dominance coefficients is effectively equivalent across ploidies.

### Ploidy effects on positive and linked selection

It has been proposed that greater mutational opportunity in tetraploids should lead to higher rates of adaptation under certain dominance conditions [16]. Using DFE-alpha analysis [38], we estimated the proportion of nonsynonymous (0-dg) sites fixed by positive selection in each population. Using either *α* or *ωα*, this proportion was significantly higher in tetraploid populations (Fig. 2E, S8 and Table S7) indicating a higher rate of adaptive substitution. This does not simply reflect the influx of introgressed alleles (below), as the difference remained significant when we removed the two tetraploid lineages admixed by diploids (*S. Carp.-4x* and *Ruderal-4x*; Table S7).

Although we find a higher proportion of adaptive substitutions, the fixation of particular mutations is generally expected to take longer in tetraploids [19], which has implications for the degree that linked selection reduces diversity during selective sweeps. We thus approximated linkage disequilibrium (LD) using the average squared genotypic correlation between SNPs as a proxy (Fig.3A). Genotypic correlations were overall significantly reduced in tetraploids, with 1kb correlations on average 50% higher in diploids than in tetraploids. We then assessed the impact on linked selection by analysing the relationship between excess nonsynonymous divergence (E_NS_ = d_N_ - d_S_) and 4-dg site diversity across genomic windows (Fig. 3B; Table S8). Regardless of ploidy, we found a consistently negative relationship between E_NS_ and 4-dg *θ_π_*, suggesting that regions of the genome that have undergone divergent selection exhibit reduced diversity at linked, neutral sites. Furthermore, we observe a highly significant quadratic effect of E_NS_, indicating that diversity is dampened for highly negative regions (i.e. those under purifying selection) as well as positive regions (i.e. sweep/divergently selected regions). The reductive effect of E_NS_ on neutral diversity was significantly stronger in gene-dense regions (upper 20%, Fig. 3D) than in gene poor regions (lower 20%, Fig. 3C). Overall neutral diversity was also significantly reduced in gene-dense regions even for windows evolving locally neutrally (E_NS_=0), consistent with an impact of linked selection acting on nearby windows rich in genes. Within these gene-dense regions, we observed significantly higher neutral diversity in tetraploids across E_NS_ values (Fig. 3D), while there was no difference in low gene-density regions (Fig. 3C), as indicated by a significant 3-way interaction between 4-dg diversity, gene-density (GDM), and ploidy. In addition, there was a difference between ploidies in the average slope in gene-dense regions but not in gene-poor regions (Fig 3D), which could suggest that linked selection in tetraploids has a relatively stronger effect within highly positively selected regions (E_NS_>>0).

**Fig 3:**
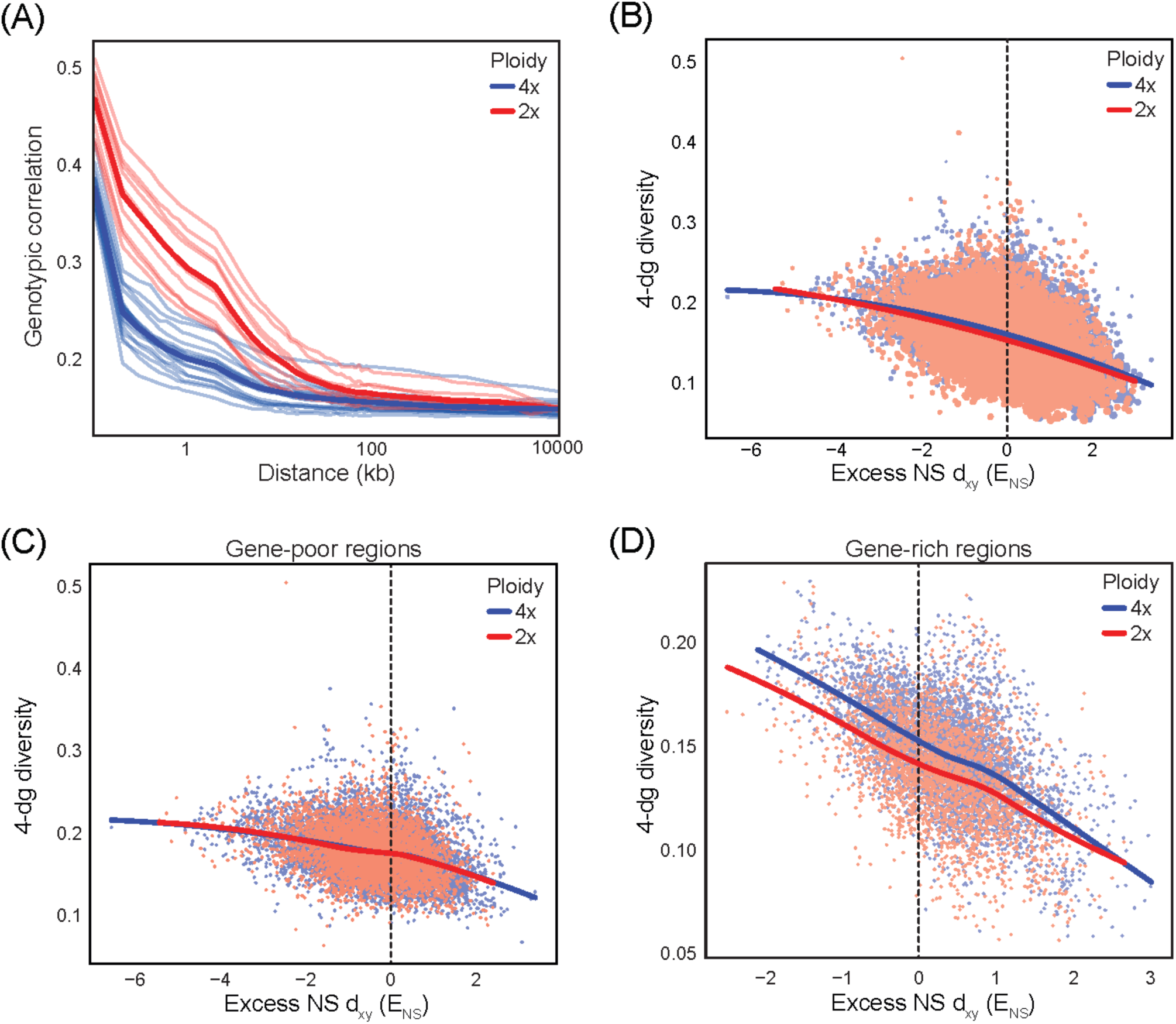
Ploidy effects on linkage disequilibrium and the strength of linked selection. (A) Decay of genotypic correlations (∼ linked disequilibrium) within each population and averaged for each ploidy (heavy lines) as a function of distance between sites (B) Curvilinear relationship between excess 0-dg d_XY_ on neutral diversity (4-dg *θ_π_*,) indicating linked selection. Size of points indicates gene density. (C-D) Linear relationship between excess 0-dg d_XY_ on neutral diversity (4-dg *θ_π_*,) for gene-poor (<20^th^ GDM percentile) and gene-dense regions (>90^th^ GDM percentile), respectively.

While slower fixation times in tetraploids would dampen a signature of linked selection, the evolution of reduced recombination in tetraploids (to avoid the formation of deleterious multivalents during meiosis [54]) as well as systematic differences across ploidies in the age of selective sweeps (due to the comparatively recent tetraploid formation) could effectively counter this effect. Such reduced recombination is not evident, genome-wide, in tetraploids. In fact, our LD approximation is generally lower in tetraploids, likely reflective of a higher population recombination rate *ρ*=8*N*_*e*_*r* (a function of effective population size and recombination rate) and/or the more recent population expansion [39]. Unfortunately, the lack of genetic maps as well as a workable phasing algorithm prevents inclusion of the recombination landscape in our regression modelling approach. Furthermore, estimation of the age and strength of selection is not currently possible on a genomic scale. Understanding the interplay between fixation times, evolution of recombination landscapes, and natural history will be the focus of future investigations.

### Single origin of tetraploids and interploidy introgression

Although previous work supported a single origin of tetraploids in the W. Carpathians [31], it is striking that in two parallel cases (Southern Carpathians and Baltic coast), local tetraploids clustered genetically with locally co-occurring diploids (Fig. 1, S3, S4B). Such a pattern suggests the possibility of multiple tetraploid origins followed by widespread gene flow among tetraploids, as these two tetraploid lineages still share a sizeable portion of polymorphisms with the widespread tetraploid lineages (*W. Carp.-4x, C. Europe-4x*, Fig. 1B). However, we find multiple lines of evidence supporting a single tetraploid origin followed by rampant local interploidy gene flow (see also Supplementary Text 1 for detailed discussion). First, coalescent simulations of population quartets involving populations from all tetraploid lineages consistently favour scenarios with a single tetraploid ancestor (∼20k – 31k generations ago) followed by admixture (Fig. 4A, 4B, S9, S10; Table S9). Second, frequencies of alleles diagnostic of the putative diploid ancestor of all tetraploids (*W. Carp.-2x* lineage) are elevated and positively correlated across all tetraploid populations (Fig. 4C, S11). Finally, alleles of several key meiosis genes are consistently shared among all tetraploids but are divergent from diploids both in and off the contact zones (Fig. 4D, S12). The extent to which interploidy gene flow can obscure the signals of tetraploid origin is an important caveat for other studies of polyploid origin and highlights the importance of considering interploidy gene flow in analyses.

**Fig 4:**
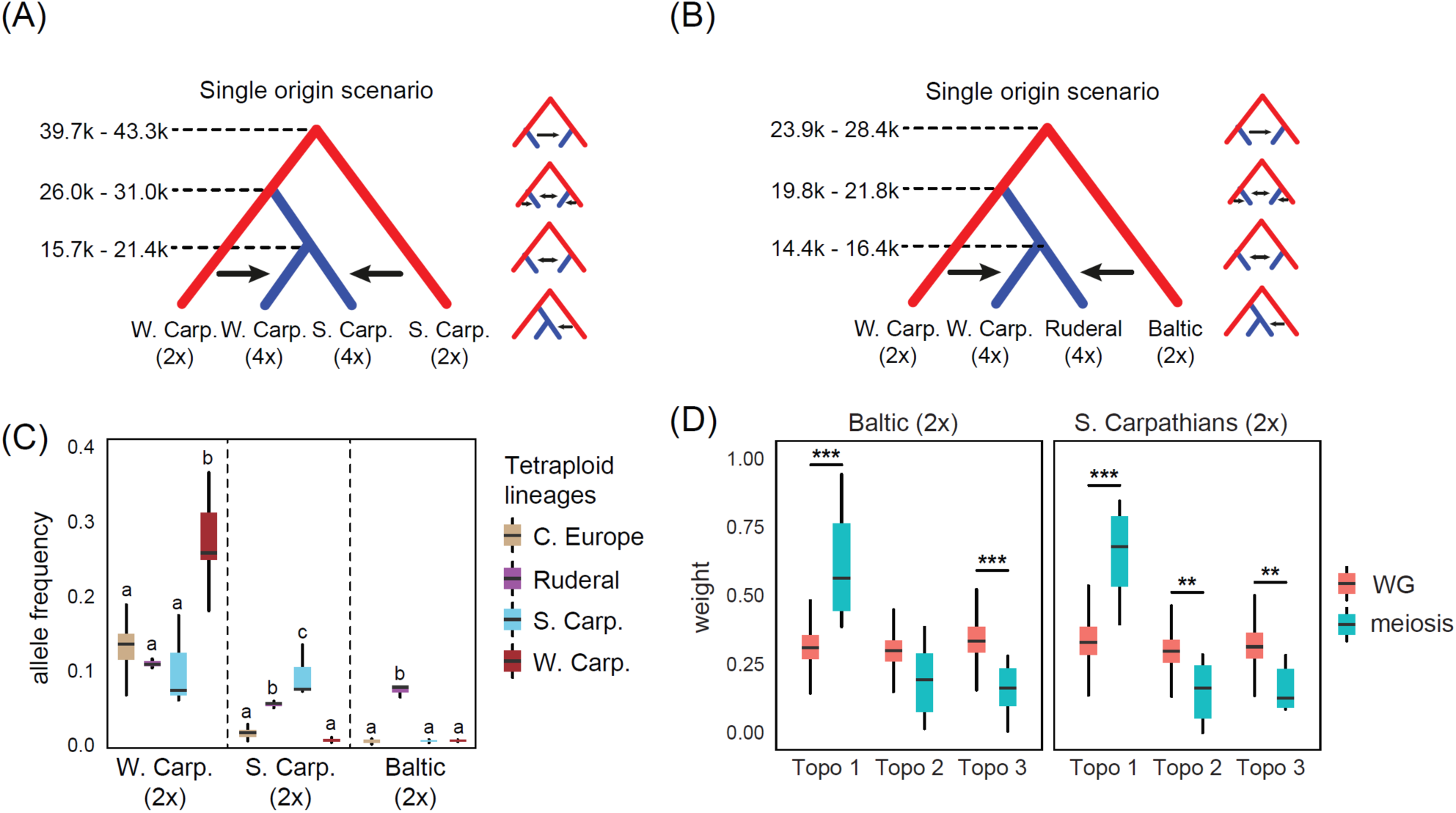
Specific and substantial admixture of locally co-occurring diploids to tetraploids. Most likely scenario inferred by coalescent simulations indicating single origin of the (A) *S. Carp.-4x*, and (B) *Ruderal-4x* tetraploids followed by local admixture from their geographically proximal diploids (large scheme) vs. representatives of the competing scenarios (smaller schemes). Median ML estimates of divergence times (range across different population quartets) in generations are above the corresponding branches. (C) Average frequencies of alleles diagnostic to particular diploid *A. arenosa* lineages present in each tetraploid lineage. Significant differences within each category of diploid alleles are designated by letters. (D) Significant prevalence of tetraploids-as-sisters topology (Topo 1) weight in set of 6 meiosis-related genes as compared with genome-wide average (WG).

In addition, multiple tetraploid (but no diploid) populations showed elevated frequencies of nuclear (Fig. S13) and occasionally also plastome (Fig. S14) markers otherwise private to *Arabidopsis lyrata*– a partially sympatric species that is known to hybridize with *A. arenosa* at the tetraploid but not diploid level [28, 40]. These admixed tetraploid *A. arenosa* populations come either from areas spatially proximal to a known hybrid zone with *A. lyrata* ([40]; some members of the *C. Europe-4x* group) or from the widespread *Ruderal-4x* lineage (Fig. S13).

The maintenance of tetraploid alleles at key meiosis genes in the face of rampant gene flow from diploids raises a question whether some regions were more prone to be locally admixed with diploid alleles while others were more resistant to introgression. To identify such regions in both the Southern Carpathian and Baltic-Ruderal contact zones, we evaluated across the genome the local weights of topologies supporting tetraploid monophyly vs. local admixture (Fig. 5, S15). In both contact zones genome-wide patterns were a complex mosaic of tree topologies, but we found localized genomic evidence of interploidy introgression from diploids into tetraploids (evidenced by greatly reduced divergence at the locus to the sympatric diploid, see Fig. 5) as well as opposite cases of genomic regions that are resistant to admixture (evidenced by strongly localised tetraploid monophyly at the locus). We also detected signals of introgression of plastid DNA in the Southern Carpathian contact zone: 37% of the *S. Carp.-4x* possessed the regionally-specific haplotypes typical for the *S. Carp.-2x*, while this pattern was absent in the *Ruderal-4x* populations which only shared plastid haplotypes with other tetraploid groups (*W. Carp.-4x* and *C. Europe-4x*).

**Fig 5:**
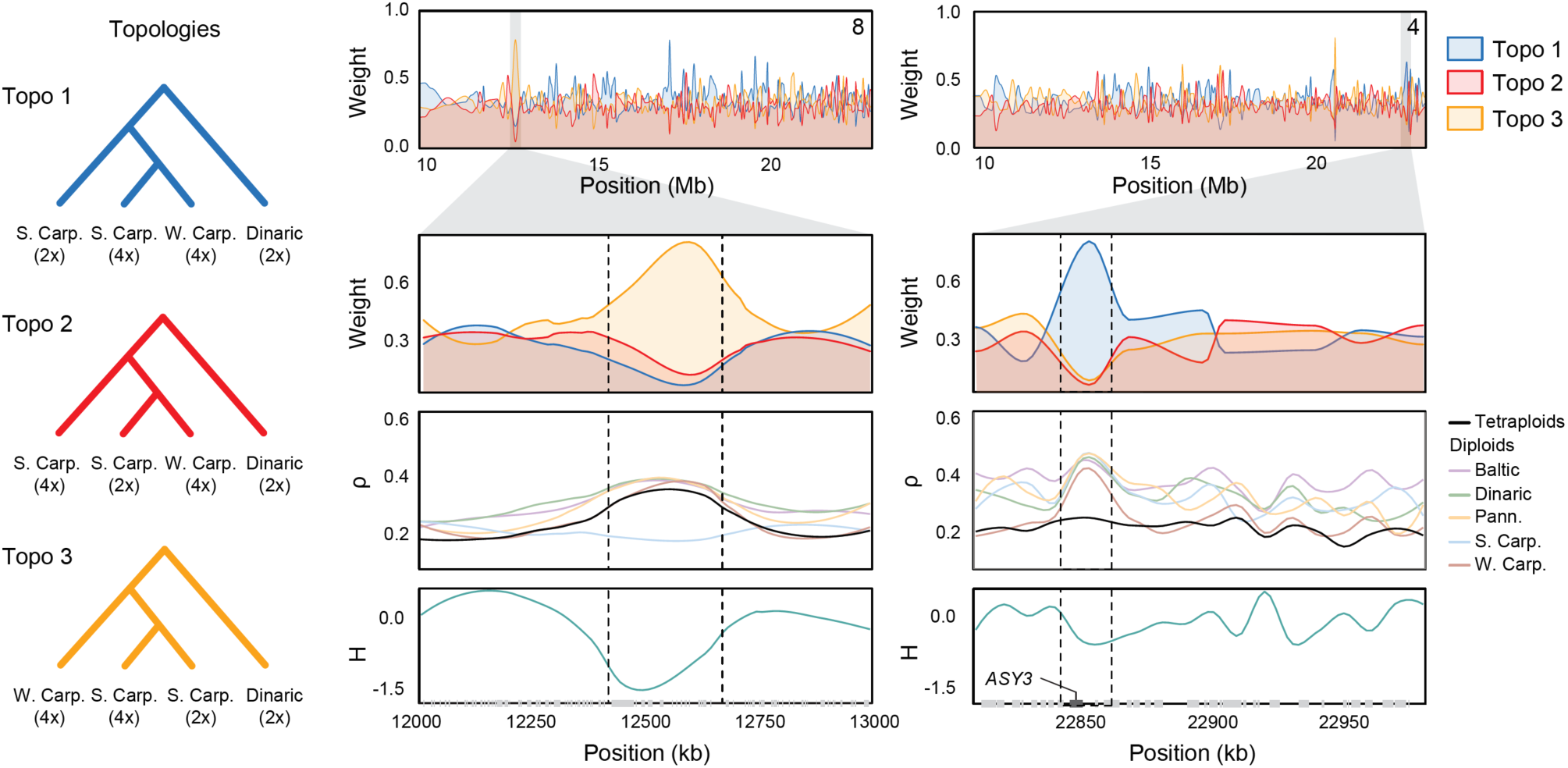
Signals of interploidy introgression and barrier loci. Topology weightings for the three diagnostic topologies relating *S. Carp.-2x, S. Carp.-4x, W. Carp-4x* and the outgroup, *Dinaric-2x.* The left zoomed-in panel represents an interploidy introgressed locus (dominant Topo 3) and positively selected (deeply negative H) region. The right panel demonstrates that a meiotic locus, ASY3, is strongly resistant to introgression (dominant Topo 1). Note that the greater breadth of all peaks relative to the meiotic locus (adjacent in figure) is consistent with much more recent introgression of these regions. From top to bottom, the zoomed-in panels depict: weight of the topology depicted on the left margin, average divergence (ρ) of the focal tetraploid relative to all other tetraploids (black line) as well as the diploid lineages, and Fay and Wu’s H.

We then tested whether positive selection was associated with these local patterns of gene flow by looking for regions that showed both elevated evidence of local admixture (Topology 3) as well as decreased Fay and Wu’s H (evidence of directional selection based on excesses of high-frequency derived alleles). In each contact zone we identified regions introgressed from diploids that also show strong marks of recent positive selection (Fig. 5; see Fig. S16 for additional cases). By overlapping 1% outliers for both metrics, we find regions containing gene coding loci (Table S10) as well as some indication of functional enrichment (see Supplementary Text 2). We also find cases of the opposite pattern, outliers for single tetraploid origin (Topology 1) and decreased Fay and Wu’s H, which suggests that diploid alleles are selected against at these loci. Consistent with a strong tetraploid resistance to diploid introgression in these regions, most of them included genes previously identified as exhibiting the strongest signatures of tetraploid-specific selection in a subset of *A. arenosa* populations previously sequenced [54]. Importantly, that these regions are resistant to introgression suggests they have an ongoing role in the maintenance of stable autopolyploid chromosome segregation and were not important only in the early stages following whole genome duplication.

## Discussion

Using the largest population resequencing dataset to date of a ploidy variable plant species, we observe pervasive differences in how forces governing genome evolution shape genetic diversity and divergence in nature. In diploid and autotetraploid *A. arenosa*, we find subtly distinct signatures of linked as well as purifying selection. Additionally, multiple sources of evidence indicate substantial gene flow from diploids to tetraploids. We discuss these results in terms of the inherent effects of the doubling of chromosomes and the possible implications for the evolutionary potential of polyploid lineages.

The effects of genome doubling on selective processes are multifarious and sometimes counter-acting, making it difficult to observe and distinguish individual causes. Additionally, signals of selection are heavily confounded with the demographic events associated with the creation, establishment, and expansion of newly formed tetraploids. For instance, although recent expansion of tetraploids reduces linkage disequilibrium globally (Fig 3A.; [39]), it can increase the apparent strength of genetic hitchhiking as strong selective sweeps in tetraploids will have had less time to re-accrue neutral diversity.

Despite these challenges, our results support the notion that both altered dominance relationship along with higher mutational input are key components of genome evolution in tetraploids. Given the strength of purifying selection is similar across ploidies (Fig. 2D), the higher ratio of (non)synonymous polymorphisms as well as the greater nonsynonymous polymorphism in highly expressed genes held under purifying selection (Fig 2A) reflect the added masking or shielding of recessive deleterious mutations in tetraploids. However, the reduced ability to see a mutation’s effect in a population is not sufficient to slow adaptation due to the fixation of beneficial alleles. The higher proportion of nonsynonymous polymorphisms fixed by positive selection in tetraploids (Fig. 2E) suggests that the increased mutational input is sufficient to overcome any hindrance to adaptation posed by the reduced efficiency of selection [16, 41], such that autotetraploid populations may actually respond more rapidly to directional selection [20]. Indeed, *A. arenosa* tetraploids expanded well-beyond their ancestor diploid’s range, including postglacial landscapes and new man-made habitats [34], suggesting an enhanced ability to establish in novel habitats. Increased mutational input and retention of diversity may be particularly apt for the fluctuating environments that are commonly seen as characteristic of polyploids [7, 41-43].

Although increased diversity is generally viewed as a driver of polyploid success, it can also be detrimental. For loci held under purifying selection, higher equilibrium values of nonsynonymous diversity are expected to result in increased genetic load [18]. However, our load analysis does not support higher load for current tetraploids, even though nonsynonymous diversity is indeed higher for genes under purifying selection. This may reflect that the younger tetraploids have simply not yet reached equilibrium, which could take hundreds of thousands of generations to establish [16]. In addition, double reduction, a unique phenomenon in autopolyploids in which the resolution of multivalents occasionally causes sister chromatids to segregate into the same gamete, may also play a role. This process will increase homozygosity and allow more efficient purging of deleterious alleles [44].

Despite an increased recognition of adaptive introgression [45], introgression from divergent lineages, species, or ploidy cytotypes may also impart maladaptive diversity [45, 46]. The most salient example for this lies in meiotic genes, which have been shown to exhibit the strongest signatures of selection in tetraploids and have evolved to ensure faithful segregation of chromosomes and proper formation of gametes [43]. The introduction of diploid-like meiotic alleles into a tetraploid population would increase the frequency of multivalent formation, and thus decrease fitness. Consistently, meiotic genes frequently show the strongest signatures of resistance to introgression in our *A. arenosa* dataset: elevated divergence between ploidies, reduced diversity within tetraploids, and tetraploid monophyly in both diploid-tetraploid contact zones (Figs. 4D, 5). Although we are not able to generally quantify its precise benefit/detriment, such evidence of resistance to introgression is strongly indicative of maladaptive gene flow. On the other hand, we also found coding regions with diploid-like derived alleles that have swept to higher frequencies in co-occurring tetraploids (Fig. 5), suggesting that in such cases the introgressed allele may be adaptive. In fact, the most widespread tetraploid lineage (*Ruderal-4x*) is the only lineage with traces of introgression from not only a distinct diploid *A. arenosa* (*Baltic-2x*) but also from a different species – *A. lyrata* (Figs. 1, S13). *Ruderal* tetraploids have coincidentally switched to a very different, weedy, life strategy [48], colonizing man-made habitats in vast areas of central and northern Europe [49]. Overall, this points to the general ability of tetraploids to accumulate diversity from various lineages, while retaining critical tetraploid- or locality-specific adaptations.

In conclusion, despite the ubiquity of polyploidy especially throughout the plant kingdom (e.g. [2, 50]), the drivers of successful establishment and spread of newly formed polyploid lineages remain obscure. In contrast to frequently studied ecological explanations [51, 52], differences in population genomic processes have not been assessed at the genomic level in natural populations despite being repeatedly invoked as potential drivers [7, 16]. Our results thus provide the first empirical insight into genomic drivers of evolutionary potential of autopolyploids. Despite slightly increased nonsynonymous diversity, tetraploids may still benefit from masking from potentially deleterious recessive mutations, and also exhibit consistently higher frequencies of adaptive nonsynonymous substitutions. Finally, multiple events of strong introgression into tetraploids may provide additional substrate for local adaptation. This supports the view of polyploids as diverse and adaptable evolutionary amalgamates from multiple distinct ancestral lineages [53].

## Online Methods

### Plant Material and Library Preparation

In addition to eight previously sequenced populations [54-56] we collected 31 new populations throughout the distribution range of *A. arenosa* (see Table S11 and Fig. S17) and its closest relative, *A. croatica*. We aimed to cover each main evolutionary lineage distinguished by previous RADseq studies [31, 34] by multiple populations and also representatively cover the ploidy level (15 diploid, 24 tetraploid populations), altitudinal (range 1 – 2,240 m a.s.l.) and edaphic variation (17 calcareous, 21 siliceous, 1 serpentine substrate).

We extracted DNA from silica-dried leaf tissue according to a CTAB protocol [57] with the following modifications: 75 – 100 mg of dry leaf tissue were ground in 2 mL tubes (Retsch swing mill), 200 units of RNase A per extraction were added to the isolation buffer and the DNA pellets were washed twice with 70% ethanol. DNA was resuspended in 50 μL TE-buffer for storage and small fragments were removed using Agencourt AMPure XP beads (Beckman Coulter, Massachusetts, USA) following the manufacturer’s instructions with 0.4x DNA:beads ratio.

We quantified the extracted gDNA using the dsDNA HS assay (Q32854) from ThermoFisher Scientific (Life Technologies Ltd. Paisley, UK) with their Qubit 2.0 or 3.0 (Q33216). We prepared Illumina (Illumina United, Fulbourn, UK) Nextera XT (FC-131-1024) and TruSeq PCR free (FC-121-3003) sequencing libraries for 350 bp insert length of genomic DNA. For PCR free libraries we used 300 to 500 ng DNA as input instead of the recommended 1 μg. We quantified the NGS libraries using Qubit as described above.

### Sequencing and Variant Calling

We multiplexed libraries based on Qubit concentration and ran those pools on an initial quantification lane. According to the yields for each sample, we increased loading of the same multiplex-mix on several lanes to achieve a minimum of 10× coverage, based on the number of raw reads. Samples that had less than our target coverage were remixed and run on another lane (top-up lane). We sequenced 125 bp pair end reads on Illumina’s HiSeq 2500 platform for all sequencing runs.

Our data processing pipeline involved three main parts: 1) Preparing the raw sequencing data, 2) Mapping and re-aligning the sequencing data and 3) Variant discovery (GATK *v.3.5*, following GATK Best Practices). All steps and parameters are summarised in File S2. To prepare the raw sequencing data for mapping we concatenated the fastq.gz files from the different sequencing lanes, followed by trimming off the adapter sequence from reads that had inserts shorter than 250 bp, using cutadapt 1.9 [58]. We mapped the reads to a North American *Arabidopsis lyrata* reference genome [59] using bwa [60]. At this stage, we added *A. arenosa* sequencing data from previous studies [6]. For Nextera (PCR-based) libraries, we removed duplicated reads using ‘MarkDuplicates’ from picard-tools 1.134 [61] followed by ‘AddOrReplaceReadGroups’ to add read groups and indices to the bam files.

We then used GATK *v.3.5* ‘RealignerTargetCreator’ and ‘IndelRealigner’ [62] to re-align the reads around indels. Prior to variant discovery, we excluded individuals that had less than 40% of bases <4× coverage (assessed via GATK ‘DepthOfCoverage’ with the restriction to a minimum base quality of 25 and a minimum mapping quality of 25). Our final dataset for analysis contained 287 *A. arenosa* and four *A. croatica* individuals from 40 populations (see File S1 for population details and File S3 for a summary of processing quality assessments).

We called variants for the 291 bam files (287 *A. arenosa* and four *A. croatica*) using ‘HaplotypeCaller’ and ‘GenotypeGVCFs’ (GATK *v.3.5*). For each bam file, ‘HaplotypeCaller’ was run in parallel for each scaffold with ploidy specified accordingly and retaining all sites (variant and non-variant). We combined the single-sample GVCF output from HaplotypeCaller to multisample GVCFs and then ran ‘GenotypeGVCFs’ to jointly genotype these GVCFs, which greatly aids in distinguishing rare variants from sequencing errors. Using GATK’s ‘SelectVariants’, we first excluded all indel and mixed sites and restricted the remaining variant sites to biallelic. Second, we removed sites that failed GATK Best Practices quality recommendations (QD < 2.0, FS > 60.0, MQ < 40.0, MQRankSum < - 12.5, ReadPosRankSum < -8.0, HaplotypeScore < 13.0). Third, we masked genes that showed excess heterozygosity (fixed heterozygous in at least five SNPs in two or more diploid populations) in the dataset, i.e. potential paralogues mapped on top of each other. At the same step, we masked sites that had excess read depth that we defined as 1.6× the second mode (with the first mode being low coverage sites indicative of mismapping) of the read depth distribution (DP > 6400).

### Polarization and Variant Classification

We repolarized a subset of sites using a collection of genotyped individuals across closely related diploid *Arabidopsis* species thus avoiding polarization against a single individual (the reference genome, N. American *A. lyrata*). We used two individuals from each of the following diploid *Arabidopsis* species (genotyped in the same way as our *A. arenosa* samples): European *A. lyrata, A. croatica*, and *A. halleri*. For a site, we considered only species with complete genotypes and only considered a site with at least two species represented. We required the alternative allele frequency to be > 0.5 in each species, if all species were represented at a site. However, if only two species were represented, we doubly weighted allele frequency for the species by preferring species with expected higher genetic variation of its European populations (i.e. with decreasing priority for *A. halleri* > *A. lyrata* > *A. croatica*) and required mean allele frequency to be > 0.5. In total, this identified ∼145,000 sites for repolarization. We classified sites as 4-fold (4-dg) or 0-fold (0-dg) degenerate based on their position in the *A. lyrata* gene model annotation Araly1_GeneModels_FilteredModels6.gff

### Population Structure

We inferred relationships among the 39 *A. arenosa* and one *A. croatica* populations (the full dataset, as well as each separate ploidy) based on putatively neutral 4-fold degenerate SNPs. Synonymous sites are not necessarily free of constraints, e.g. due to potential codon usage bias, but are nevertheless the closest to effectively neutral of any site class in the genome [63]. After quality filtering our demographic analysis is based on a genome-wide dataset consisting of 1,350,328 four-fold degenerate SNPs, allowing for a maximum of 10% missing alleles per site (1.2% missing data). Firstly, we calculated principal component analysis (PCA) using *glPCA* function in *adegenet* [64] replacing the missing values (1.2% in total) by average allele frequency for that locus. Next, we calculated Nei’s [65] distances among all individuals in *StAMPP* [66] and displayed it using the neighbour network algorithm in *SplitsTree* [67]. Third, we selected the 553 (503 for the diploid only dataset) most parsimony-informative genes based on missing data filter criteria (accessions with ≥ 10% missing data per gene were omitted from the respective gene and those genes with ≥ 10% missing accessions per gene omitted) and constructed a maximum likelihood tree from each gene using *RAxML v.8* [77] with model GTRCAT and rapid bootstrap with 100 replicates each. In each gene alignment for *RAxML*, accessions were represented by the consensus sequence, with different alleles represented as ambiguous sites in the consensus sequence. Ambiguous sites are treated by *RAxML* as invariant sites, hence, the standard nucleotide substitution model needed to be utilized; the ascertainment bias correction model that is usually used for SNP matrices is not appropriate in such case. The resulting gene trees were summarized under the multispecies coalescent using *Astral v.4.10.10* [78], bootstrapping was performed with 100 replicates each.

We further determined grouping of the populations using three clustering approaches: model-based Bayesian clustering using *fastStructure v.1.0* [68] and STRUCTURE *v.2.3.2* [69] and a non-parametric k-means clustering using *adegenet* [64]. The analyses were performed separately for (i) the entire data set of *A. arenosa* (*A. croatica* excluded; 9,543 SNPs after random thinning over windows of 50 kb to reduce effect of linkage and removing singletons, 2.4% of missing data), (ii) diploids only (12,655 SNPs, 4.1% missing data) and (iii) tetraploids only (9,596 SNPs, 2.3% missing data). In *fastStructure*, five replicate runs for K (number of groups) ranging from 1 to 10 were carried out under default settings. We selected the optimal K value based on the similarity coefficient (∼1 for optimal K [70]) across replicates (Fig. S18). As *fastStructure* does not handle polyploid genotypes, we randomly subsampled two alleles per each tetraploid locus (following [71]) using a custom script. To check for the effect of such subsampling, we also ran the original STRUCTURE program, which handles mixed-ploidy datasets, for optimal K values according to *fastStructure*. We ran the admixture model with uncorrelated allele frequencies using a burn-in of 100,000 iterations followed by 1,000,000 additional iterations. Finally, we ran k-means clustering using 1000 random starts and selected the partition with the lowest Bayesian information criterion (BIC) value.

We used Treemix *v.1.3* to infer migration events and relationships between the 39 *A. arenosa* populations using one *A. croatica* population as outgroup. We used the 4-dg sites to build a tree without any migration events and used this tree as basis for migration models to make comparisons easier (option ‘-g’). We modelled no to eight migrations and graphically assessed the residuals after each additional migration modelled, using the R-scripts supplied with the Treemix package. If specific population pairs had high residuals, we modelled an additional migration event. We continued until the residuals were small and evenly spread across population pairs and/or until an additional migration event involved the outgroup (we consider this admixture unlikely due to very local occurrence and spatial isolation of the *A. croatica*).

To quantify differentiation among populations, we calculated genome-wide F_ST_ and Rho coefficients (similarly as in the window based analyses described below) and performed analysis of molecular variance (AMOVA) based on the Nei’s distances using the *amova* function in the *pegas* R package [72]. We tested for isolation by distance relationships through comparison of matrices of geographic and genetic (Nei’s among-population) distances among the populations using *mantel.randtest* function in *ade4* R package [73]. For each tetraploid population, we calculated the frequency of alleles diagnostic to each diploid lineage. The allele was defined as diagnostic if it exhibited min. frequency 0.3 in that diploid lineage and was absent in any other diploid lineage (except for the putatively admixed Baltic diploids, Table S13). For all populations we also calculated frequency of *A. lyrata*-like alleles, i.e. reference alleles that were otherwise rare in the complete *A. arenosa* dataset (a rarity cut-off of 6.8%, i.e., equivalent to two tetraploid populations of 8 individuals). As these alleles were nearly absent in *A. arenosa* diploid populations, i.e. the ancestors of tetraploids, we assume they more likely represent hybridisation from *A. lyrata* than ancestral variation shared among both species.

Finally, we inferred phylogenetic relationships among plastomes of our samples and previously published plastomes of other *Arabidopsis* species [71]. We mapped the reads to a custom *A. arenosa* plastome assembly constructed using org.ASM (http://pythonhosted.org/ORG.asm/) and performed variant calling and filtration as described above, with the exception of setting ploidy = 1 in GATK *HaplotypeCaller* and retaining SNPs and invariant sites with depth > 4 in at least 90% of the individuals. We aligned all sequences using *Mafft* [74] and reconstructed relationships using maximum likelihood in *RAxML* using GTR model with Gamma distribution of rate variation.

### Demographic analysis

We compared various demographic models and estimated parameters using the coalescent simulation software *fastsimcoal2 v.25* [75]. The models differed in topology and presence/absence of migration (admixture) events (Figs 4, S9, and S10), and each model was fit to a multi-dimensional site frequency spectrum calculated from the observed four-fold degenerate SNP data. Our primary interest in these analyses lie in confirming whether or not the additional populations that we sampled supported the single origin of tetraploids previously determined in [31]. Specifically, we focused on populations in the two diploid/tetraploid contact zones (Southern Carpathian and Baltic-Ruderal contact zones).

We attempted to discriminate between single versus independent origins using population quartets involving representatives from both putative parental diploid lineages (*S. Carp-2x* and *W. Carp.-2x* for *S. Carp.-4x*; *Baltic-2x* and *W. Carp.-2x* for *Ruderal-4x*; i.e. the closest two in the descriptive analyses, Fig. 1 and 4), the *W. Carp.-4x* that is closest to the putative ancestor of the widespread tetraploids [31] and the focal tetraploid (Fig. S9 and S10). In order to maintain a realistic number of scenarios while permuting the parameters (11 models for each population quartet), we modelled both uni- and bi-directional admixture within the same ploidy level, but only unidirectional interploidy admixture – from diploids to tetraploids. This decision reflects no signs of admixture of the diploids in clustering analyses (in contrast to the highly admixed tetraploids, Fig 1B) and virtual absence of triploids in nature [32], i.e. the only possible mediators of gene flow in the tetraploid-to-diploid direction [52]. In addition, we tested for the potentially admixed origin of the Baltic diploids (*Baltic-2x*) [34] using population trios involving representatives of each diploid lineage (*W. Carp.-2x* and *S. Carp.-2x*) as well as the focal *Baltic-2x* population (Fig S19 and Table S4).

For each scenario and population trio/quartet, we performed 50 independent *fastsimcoal* runs to overcome local maxima in the likelihood surface (see File S7 for example template file). In order to minimize the population-specific effects, we ran the analyses for different iterations of well-covered populations falling within the particular lineage, leading to 12 different population quartets (“natural replicates”) for each scenario testing the origin of the *S. Carp.-4x* and *Ruderal-4x* and four trios in the *Baltic-2x* scenarios. We then extracted the best likelihood partition for each *fastsimcoal* run, calculated Akaike information criterion (AIC) and summarized them across the 50 different runs, over the scenarios and different population trios/quartets. The scenario with consistently lowest AIC values within and across particular population trios/quartets was preferred (Figs. S9 and S10). In order to get confidence intervals for the demographic parameters (Table S9), we sampled with replacement from the 4-dg SNPs to create 100 bootstrapped datasets and performed additional *fastsimcoal2* analyses under the preferred scenario with these 100 distinct datasets. For these analyses we also included representative (best covered) populations from the putatively non-admixed *C. Europe-4x* lineage. Finally, we used the mutation rate of 4.3×10^-8^ estimated by [31] to calibrate coalescent simulations and obtain absolute values of population sizes and divergence times.

In addition, we used PSMC 0.6.4 [76] to infer changes in effective population size (N_e_) through time using information from whole-genome sequences of the *A. arenosa* diploids. We plotted 75 samples out of the 93 sequenced diploids, i.e. excluding samples with too low a coverage (below 12×) and too much missing data. Coverage and missing data might have large effects to the PSMC estimates [77] and hence our results should be interpreted only in conjunction with other analysis methods. We run PSMC with parameters: psmc -N25 -t15 -r5 -p “4+25*2+4+6” and then plotted the past changes in N_e_ assuming a mutation rate of 3.7×10^-8^ substitutions per site per generation and generation time of two years.

### Window-based metric calculation

In order to facilitate comparisons of windows across populations or population contrasts, we chose to calculate population genetic metrics in windows defined by a given number of base pairs. We repeated all calculations for two window sizes, 10kb and 50kb. We used the 50kb windows for characterizing broad, genome or chromosome-level patterns, whereas the former was used for finer, gene-level analyses. For 50kb windows, patterns of LD decay suggest a minimal degree of non-independence among windows relative to the genome background (Fig. 3A).

For each of the 36 populations with at least five individuals, we excluded all individuals with < 8× average coverage, except for populations SZI, KZL, and SNO as excluding individuals from these populations would drop them below required minimum of 5 individuals. After excluding these individuals, we excluded sites if the number of missing individuals was greater than 10%, on a population specific basis. When calculating diversity, we downsampled each population to 5 individuals on a per-site basis. We calculated diversity as θ_π_ [78] divided by the total number of sites with sufficient coverage.

We calculated the following divergence metrics for each possible pairwise population comparison using our custom scripts available at https://github.com/pmonnahan/ScanTools: F_ST_ [79], ρ [17], d_XY_ [80], and the number and proportion of fixed differences. The multi-locus implementation of F_ST_ and ρ was translated from the software SPAGeDi [81].

### Topology weighting and detection of local introgression

We quantified the relative support for alternative phylogenetic relationships among populations using the topology weighting approach implemented in Twisst [82]. We used only 4-fold degenerate sites and used only individuals with > 8x coverage. Using bcftools, we converted the VCF files to a simplified tabular genotype file containing only the relevant individuals. We filtered this file using the filterGenotypes.py script that accompanies the Twisst software. At a site, we required genotype calls for at least 200 out of the 254 high coverage individuals (i.e. allowing ∼20% missing data). We used only biallelic sites and required that the minor allele be present in at least 2 individuals. We then ran phyml_sliding_windows.py using 100 SNP windows (-w 100 and –M 20), which fits an ML phylogenetic tree for each window. Ideally, Twisst should be run on phased data; however, we were unable to find a workable phasing software that could handle diploids and tetraploids despite multiple attempts. Instead, we used the phasing algorithm internal to Twisst, which forms haplotypes by maximizing pairwise LD in each window.

We then ran Twisst for a number of scenarios, specifying individual population or groups of populations (lineages) as taxa. Twisst implements an iterative sub-sampling algorithm based on the phyML results to determine the support or weight of each possible taxon topology within each window. We requested the program calculate the complete weightings (completely searching sample space) if possible and used an approximate method, where sampling ceases after a given threshold of confidence is reached, when necessary. We allowed for 2000 sampling iterations before opting for the backup method. After this limit, we used the “Wilson” method at the 5% level, which will enforce sampling until the binomial 95% confidence interval is less than 5% of the weight value.

We used a combination of information from Twisst as well as divergence metrics to diagnose regions of both excessively strong and weak interploidy introgression in the two highly admixed *S. Carp.-4x* and *Ruderal-4x* lineages. First, introgressed regions should show an elevated weight for topologies wherein the proximal diploid/tetraploid pair are placed sister to one another (Topology 3 in Fig. 5). Second, when comparing the focal tetraploid to other tetraploid populations, an introgressed region should show elevated divergence while at the same time exhibiting reduced divergence to the focal diploid population. Conversely, introgression-resistant regions should show elevated Topology 1 and a combination of low divergence from tetraploids and elevated divergence from all diploids. We looked for evidence of selection on introgressed regions by overlapping window outliers for Topology 3 and Fay and Wu’s H (in 10kb windows) in the focal tetraploid (99^th^ percentile for both metrics).

### Gene expression analysis of purifying selection

We evaluated patterns of diversity at the gene level using gene expression levels as a proxy for selective pressure based on evidence that higher-expressed genes generally show stronger signs of purifying selection in both plants and animals [35, 83-85]. To obtain gene-wise estimates of diversity, we performed a separate mapping process (again, using *A. lyrata* as the reference genome) using a subset of the total *A. arenosa* dataset that covers all major diploid and tetraploid lineages (9 tetraploid and 9 diploid populations, comprising 74 and 70 individuals, respectively, listed in Table S12). We retained sites with read depth of 4 or higher for at least 5 individuals across each population (9 – 14 million sites per population, Table S12). Sites with more than 5 individuals covered were down-sampled to 5 to homogenize chromosome depth across sites.

First, we extracted RNA from leaves of 3-week old individuals with three biological replicates for each of three diploid populations (HNI, RZA, SNO) to complete our previous dataset [86] of seven tetraploid populations (TBG, BGS, STE, KAS, CA2, HOC, SWA) using the RNeasy Plant Mini Kit (Qiagen). We synthesized single strand cDNA from 500ng of total RNA using VN-anchored poly-T(23) primers with MuLV Reverse Transcriptase (Enzymatics) according to the manufacturer’s recommendations. We made RNAseq libraries using the TruSeq RNA Sample Prep Kit v2 (Illumina) and sequenced libraries on an Illumina HiSeq 2000 with 50bp single-end reads. We sequenced between 9.8 and 18.8 million reads (avg 13.6 million). We aligned reads to the *A. lyrata* genome using TopHat2 [87] and re-aligned unmapped reads using Stampy [88]. We acquired read counts for each of the 32,670 genes using HTseq-count [89] with *A. lyrata* gene models. We normalized for sequencing depth using DEseq2 in R [90] and further analyses were performed in MATLAB (MathWorks).

Analysis of differential expression between diploid and tetraploid expression patterns were performed using a one-way analysis of variance (ANOVA), and *p*-values were corrected for false discovery rate [91]. To avoid low-expression genes, we filtered for genes presenting a least one sample with normalized counts above 25, and computed the log-ratio of the average population expression in tetraploid populations against the average expression in diploids (positive when the expression of a gene is higher in tetraploid and negative when it is higher in diploids).

We obtained 6,504 genes with statistically significant differential expression (*p* < 0.05) between diploids and tetraploids (33% of 19,319 genes), but only 321 of these presented fold-change above 1.78x (5% two-tail threshold, Fig. S20A) and 214 above 2x. Overall, the average mean expression across populations is very strongly correlated between ploidies (slope = 1.02, R^2^ = 0.93, Fig. S20B), and to estimate mutational patterns we limited ourselves to the set of 18,998 genes non-differentially expressed (NDE) between ploidies.

We then filtered genes for independence of diversity metrics from number of sites, specifically those that showed a correlation of number of sites with diversity (indicating potential mis-mapping of reads; Fig. S21). This effect of 4-dg *θ_π_* and *θ_W_* was strong for genes with less than 20 sites or more than a 100 using a locally weighted linear regression (LOWESS) for genes with a minimum of 5 sites of each fold (0-dg and 4-dg). Between these two boundaries, the number of sites only has a weak effect on 4-dg diversity. We observed a similar pattern in terms of 0-dg diversity with loci with less than 30 or more than 400 0-dg sites (Fig. S21 C&D). After exclusion of loci outside of these bounds (for both 4-dg and 0-dg) from any downstream analysis we were able to cover around 45% of all NDE genes.

We then visualized the correlation of diversity of each gene with the average gene expression within the ploidy of the population with a locally weighted linear regression (LOWESS). For genes with expression levels above a certain expression threshold (50), nonsynonymous diversity (0-dg *θ_π_* and *θ*_*W*_) showed a clear negative correlation with expression (proxy for strength of purifying selection) for both ploidies (Fig. 2A, Fig. S6: bold lines). Notably, this trend seems to break for very high expressions (>2250 i.e. top 0.35%) possibly due to the low coverage of this expression range (67 genes). After removal of these genes outside of these thresholds, we obtained 5,900 NDE genes per population to be used for multiple linear model (MLM) fitting.

We evaluated the effect of gene expression on 0-dg/4-dg diversity ratio for each population by modelling it as a function of its ploidy (p) with coefficient α_p_, the average gene expression measured in ploidy p (E_p_) with coefficient β, and an interaction term γ_p_ as follows:

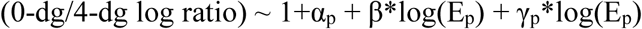

The second MLM equation for evaluating the impact of population size on 0-dg diversity was established as follows using stepwise regression evaluating each term based on the *p*-value for an *F*-test of the change in the sum of squared error by adding or removing the term.

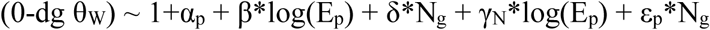

where the interaction term with log expression γ is now dependent on N_g_, δ represents the fixed effect of N_g_, with an additional interaction term εp dependent on ploidy (p). To estimate N_g_, we first estimated effective population sizes using synonymous diversity as an estimator of θ, the estimated mutation rate (μ) of 4.3×10^-8^ for *A. arenosa* [31] and their theoretical relationship given by θ=4μN_e_ in diploids and θ=8μN_e_ in tetraploids. This gave an estimate of effective population sizes around 240,000 individuals for diploids and around 130,000 for tetraploids. In terms of number of haploid genomes, this difference in effective census sizes is more than compensated by tetrasomy (∼480,000 in tetraploids vs ∼520,000 in diploids). The MLM estimates are presented in Table S5 and S6, and the estimated effects for values of the predictor chosen to show large responses are plotted in Fig 2B: log Expression: 3.9124 to 7.7098; Ng: 366058 (low) to 488976 (med) to 611894 (high).

In addition, we calculated recessive load as a number of sites with derived allele in homozygote state per each individual with at least 5 million SNPs called (240 individuals in total) and tested for difference among population means of diploid and tetraploid populations using Wilcoxon rank sum test.

### Distribution of fitness effect

Using the allele frequency spectra (AFS) for 4-dg and 0-dg sites (separately) for each of the 36 populations with ≥ 5 individuals screened, we estimated the distribution of fitness effects (DFE) [38], the proportion of adaptive substitutions relative to the total number of nonsynonymous substitutions (*α*) [92], the proportion of adaptive substitutions relative to neutral divergence (*ω*_*a*_; [93]). For all parameters estimated, we obtained 95% confidence intervals by analyses of 200 bootstrapped data sets. For each population, we fit two demographic models (constant population size and stepwise population size change), selected the best-fit model using a likelihood ratio test (LRT) and then estimated the parameters of the DFE, *α* and *ω*_*a*_ under this model. The DFE is estimated using a gamma distribution with a shape parameter (β) and a scale parameter that represents the strength of purifying selection. As the strength of selection is dependent of the effective population size *N*_*e*_, the result of DFE are often summarized by binning the distribution in 3 bins of *-N*_*e*_\**s*. A *-N*_*e*_*s* of 0-1 represent nearly neutral sites, 1-10 mildly deleterious and > 10 highly deleterious mutations.

For all populations, the stepwise population size change model was preferred. We ran DFE-alpha using both unfolded and folded site frequency spectra. As the results were very consistent using the folded or the unfolded allele frequency spectrum we to focus on estimates based on the folded spectra, which should be more robust. We tested whether diploids and tetraploids differed with respect to the proportion of new nonsynonymous mutations in each bin, using Wilcoxon rank sum tests.

### Linked selection analysis and calculation of genotypic associations (linkage disequilibrium)

We inspected the relationship between the excess nonsynonymous divergence (d_XY_) relative to synonymous divergence, as a proxy for divergent selection, and synonymous diversity (θ_π_) in 50kb windows [94]. Both nonsynonymous and synonymous divergence was calculated for each population in each window as the average divergence at (non)synonymous sites for all pairwise contrasts between the focal population and all other populations in the dataset. We natural-log transformed these values and standardized them to be on the same scale. Then, we simply took the difference between the scaled, transformed divergence values in each window. We refer to this difference as E_NS_. We also square root transformed θ_π_ for normality purposes, removed windows with fewer than 20 SNPs, and removed populations with fewer than 2,000 non-missing windows, retaining a total of 27 populations (10 diploid and 17 tetraploid, listed in Supplementary File S1) and an average of 2,660 windows per population (∼60% of genome). A negative relationship with θ_π_ is interpreted as evidence of a reductive effect of selection on linked, neutral diversity (i.e. linked selection). More specifically, we were interested to see if this relationship was dependent on ploidy level (i.e. is linked selection more effective in diploids or tetraploids?).

We used a multiple regression approach to infer this relationship and its dependence on ploidy level. We also included information on gene density (the proportion of bases in the window occupied by genic sequences according to the *A. lyrata* annotation) and proportion of missing data in each window. When calculating missingness in each window, we considered all biallelic sites and simply averaged the proportion of missing data across all 287 individuals in the study at each site within the window. Given the strong negative relationship between gene density and missingness, we combined them into a single, compound variable

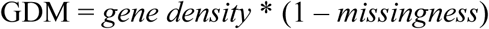

where high values indicate windows with high gene density and low missingness and low values indicate the opposite. We fit a mixed model, using the ‘lmer’ function from the R package *lme4* [95], with E_NS_ and GDM as continuous variables, ploidy as fixed categorical variable, and populations as a random categorical variable. We also included a quadratic effect of E_NS_ to investigate the possibility of a nonlinear relationship with neutral diversity. Our initial model included all possible interactions, and we selected our final model by eliminating non-significant higher order interaction terms. The results were not qualitatively different following removal of tetraploid populations admixed by non-sister diploids (*S. Carp -4x*: DRA, LAC and TZI, and *Ruderal-4x:* KOW, STE and TBG).

To calculate genotypic correlations, we recoded genotypes to represent the number of alternative alleles (0 - 2 for diploids and 0 - 4 for tetraploids). We calculated *r*^*2*^ for pairs of loci and is simply the square of the correlation coefficient. An *r* value of 1.0 (and thus an *r*^*2*^ of 1.0) means that genotypes are perfectly correlated for a particular pair of loci. However, since we do not have phase information, this *r*^*2*^ value is not equivalent to the *r*^*2*^ often reported when discussing LD. Therefore, we do not technically measure LD, but rather a related measure of genotypic associations.

To visualize LD decay (Fig. 4A), we averaged *r*^*2*^ value for all pairs of loci that fall in bins of a given distance apart. We only considered populations that have sample size of 8 or greater, and we downsampled populations with greater than 8 individuals on a per-site basis. We allowed for 1 missing individual in each population and performed the *r*^*2*^ calculation for each population separately to avoid confounding effects of population differentiation.

To observe the impacts of various factors in our data on our LD approximation, we simulated unlinked data and varied the number of sites and individuals as well as ploidy. At each site, we randomly drew allele frequencies from a uniform distribution, and then drew genotypes from the binomial distribution with *p* equal to the drawn allele frequencies and *n* of 2 or 4, depending on ploidy. The average *r*^*2*^ value for each data set indicates that the number of individuals is the primary determinant of the expected *r*^*2*^ value for unlinked sites, with the other factors exhibiting a negligible effect (Fig. S22).

## Code Availability

Custom scripts used for this study are available at https://github.com/pmonnahan/ScanTools.

## Data Availability

Sequence data that support the findings of this study have been deposited in the Sequence Read Archive (SRA; http://www.ncbi.nlm.nih.gov/sra) with the primary accession code PRJNA484107 (available upon publication of this manuscript at http://www.ncbi.nlm.nih.gov/bioproject/484107]

## Acknowledgements

The authors thank Eliška Záveská, Magdalena Lučanová and Stanislav Španiel for help with fieldwork and John Brookfield and Simon Martin for helpful comments on versions of the manuscript. LY acknowledges funding from the European Research Council (ERC) under the European Union’s Horizon 2020 research and innovation programme (grant agreement 679056) and the UK Biological and Biotechnology Research Council (BBSRC) via grant BB/P013511/1 to the John Innes Centre. KB acknowledges European Research Council Consolidator grant: CoG EVO-MEIO 681946 US National Science Foundation: IOS-1146465. Additional support was provided by Czech Science Foundation (project 16-10809S to KM), Charles University (project Primus/SCI/35 to FK), and a SNSF Early Postdoc Mobility fellowship (P2ZHP3_158773 to CS).

## Author Contributions

LY, KB, FK, PB and PM conceived the study. PM, FK, PB, BL, CS, JK, RH, RS and PP performed analyses with input from LY, KB, RH, and TS. CS, PB, GF, MB and CW performed laboratory experiments. PM, FK and PB wrote the manuscript with primary input from KB, LY, BA, CS and TS. All authors edited and approved of the final manuscript.

## Materials & Correspondence

Correspondence and material requests should be addressed to Levi Yant at.

